# Widespread brain activity increases in frontal lobe seizures with impaired consciousness

**DOI:** 10.1101/2024.09.20.614207

**Authors:** Elaheh Salardini, Aparna Vaddiparti, Avisha Kumar, Reese A. Martin, Rahiwa Z. Gebre, Christopher Andrew Arencibia, Monica B. Dhakar, Eric H. Grover, Imran H. Quraishi, Eliezer J. Sternberg, Ilena C. George, Adithya Sivaraju, Jennifer Bonito, Hitten P. Zaveri, Leah M. Gober, Shamma Ahmed, Shivani Ghoshal, Kun Wu, Pue Farooque, Lawrence J. Hirsch, Eyiyemisi C. Damisah, Jason L. Gerrard, Dennis D. Spencer, Ji Yeoun Yoo, James J. Young, Daniel Friedman, Jennifer Shum, Hal Blumenfeld

**Author notes:** Correspondence to: Hal Blumenfeld, MD, PhD, Yale Departments of Neurology, Neuroscience, Neurosurgery 333 Cedar Street, New Haven, CT 06520-8018.

## Abstract

Impaired consciousness is a serious clinical manifestation of epilepsy with negative consequences on quality of life. Little work has investigated impaired consciousness in frontal lobe seizures, a common form of focal epilepsy. In temporal lobe seizures, previous studies showed widespread cortical slow waves associated with depressed subcortical arousal and impaired consciousness. However, in frontal lobe epilepsy, it is not known whether cortical slow waves are present, or whether a very different cortical activity pattern may be related to impaired consciousness.

We used intracranial EEG recordings of 65 frontal lobe seizures in 30 patients for quantitative analysis of ictal cortical activity and its relationship to impaired consciousness. Behavioral changes based on blinded review of seizure videos were used to classify focal aware, focal impaired awareness, and focal to bilateral tonic-clonic seizures. Changes in intracranial EEG power from preictal baseline were analyzed in different cortical regions and across frequency ranges in these three categories.

We found that frontal lobe focal aware seizures showed approximately 40% increases in intracranial EEG power localized to the frontal lobe of seizure onset across frequency ranges, with relatively smaller changes in other cortical regions. Frontal lobe focal impaired awareness seizures showed approximately 50% increases in intracranial EEG power, not significantly different from focal aware seizures in the frontal lobe of seizure onset (*P* = 1.038), but significantly greater than focal aware seizures in other broad cortical regions (*P* < 0.001). Importantly, the widespread cortical increases in EEG power observed in focal impaired awareness versus focal aware seizures were seen not just in the frequency range of slow waves, but were also observed across other frequencies including fast activity. However, the widespread cortical increases in focal impaired awareness seizures differed from focal to bilateral tonic-clonic seizures where intracranial EEG power increased to a much higher level by approximately 600%. The large power increases in focal to bilateral tonic-clonic were significantly greater than in focal impaired awareness seizures both in the frontal lobe of seizure onset and in other cortical regions (*P* < 0.001).

Our findings contrast with focal temporal lobe epilepsy, where impaired consciousness is associated with cortical slow waves. We can speculate that different focal seizure types produce impaired consciousness by impacting widespread cortical regions but through different physiological mechanisms. Insights gained by studying mechanisms of impaired consciousness may be the first step towards developing novel treatments to prevent this important negative consequence of epilepsy.

## Introduction

Consciousness is essential to normal human life. Both impaired and normal consciousness have their anatomical basis in widespread regions of association cortex regulated by subcortical arousal systems.^1–5^ Impaired consciousness is a serious clinical manifestation in epilepsy, impacting quality of life, safety, and emotional health. In recent decades, several studies have investigated mechanism of impaired consciousness during generalized and focal seizures, with the largest number of investigations on temporal lobe epilepsy (TLE), the most common type of focal epilepsy.^2,6–8^ Despite all this progress, there is limited information on mechanisms leading to impaired consciousness in frontal lobe epilepsy (FLE), the second most common type of focal lobe epilepsy.

Previous studies of TLE suggest that at least two mechanisms may contribute to impaired consciousness, and these mechanisms are not mutually exclusive. One mechanism, based on the global neuronal workspace theory of consciousness, posits that impaired consciousness in TLE is associated with abnormal cortical-cortical and cortical-thalamic synchrony.^6,7,9–11^ A second mechanism, the network inhibition hypothesis, posits that focal seizures in TLE inhibit subcortical arousal systems, leading to encephalopathy-like cortical slow waves and impaired consciousness.^3,8,12,13^ Intracranial electroencephalography (icEEG) studies in TLE have shown widespread association cortical slow wave activity in the delta frequency (1 – 4 Hertz (Hz)) range outside the region of seizure onset in temporal lobe seizures associated with impaired consciousness.^8^ The network inhibition hypothesis has been further supported by extensive work in animal models demonstrating that decreased subcortical arousal leads to cortical slow waves and impaired consciousness in focal hippocampal seizures.^14–24^ Studies based on both proposed mechanisms of impaired consciousness in TLE have contributed to the development of novel neurostimulation therapies aimed at restoring consciousness during and after seizures.^9,25–29^

Focal frontal lobe epilepsy differs from TLE, as FLE exhibits greater clinical and behavioral heterogeneity ranging from paroxysmal emotional outbursts, or unilateral elementary motor signs, to large amplitude bilateral hyper-kinetic movements. Some of the electroclinical heterogeneity in FLE may be attributed to the large anatomical representation and variegated functions of the frontal lobes. Impaired consciousness in FLE can vary from full awareness and responsiveness to environmental stimuli to complete loss of consciousness.^30–35^ Investigation of the mechanisms of impaired consciousness in FLE has so far been relatively limited to just one systematic study, which demonstrated that increased synchrony between prefrontal and parietal association cortices correlates with impaired consciousness, in keeping with the global workspace theory.^36^ However, it is not known whether like in TLE, additional mechanisms may also contribute to impaired consciousness in FLE. For example, an important question is whether impaired consciousness in FLE, like TLE is accompanied by widespread cortical slow wave activity outside the region of seizure onset, or whether a different activity pattern may be present. A predominance of slow wave activity outside the region of seizure onset might suggest that impaired consciousness in FLE, like TLE, is mediated at least in part by depressed subcortical arousal leading to widespread cortical dysfunction. On the other hand, different activity patterns, aside from slow waves in widespread cortical areas, would suggest that mechanisms for impaired consciousness in FLE and TLE differ.

Therefore, our goal was to investigate the patterns of electrical activity in the human brain using icEEG signals analyzed across a range of frequencies in a relatively large sample of focal frontal lobe seizures with and without impaired consciousness. We defined focal aware (FA) and focal impaired awareness (FIA) seizures based on behavioral review. Our analysis demonstrated that frontal lobe FIA seizures had abnormal increased activity in widespread brain regions in contrast to FA seizures where increased activity was confined mainly to the frontal lobe of seizure onset. Importantly, the increased activity in frontal lobe FIA seizures was seen across a broad range of frequencies, not just slow waves, therefore differing from TLE. We next wondered if frontal lobe FIA seizures differ from focal to bilateral tonic-clonic (FBTC) seizures, which are also known to involve widespread brain areas across a range of frequencies. We found that icEEG signals did indeed increase in frontal lobe FBTC seizures across widespread brain areas and frequencies, but these increases were over 10 times larger than in frontal lobe FIA seizures.

## Materials and methods

All procedures were in accordance with the institutional review boards for human studies at Yale University School of Medicine, NYU Grossman School of Medicine, and Icahn School of Medicine at Mount Sinai. Informed consent was obtained from all participants according to the Declaration of Helsinki.

### Patients and seizures

Based on patients’ medical records, we initially identified 508 seizures from 34 consecutive patients who underwent icEEG evaluation for possible FLE between 1997 to 2020 at Yale, 2010 to 2019 at Mount Sinai, and 2010 to 2019 at NYU. Seizures and patients were excluded from the final analysis using the criteria below.

Based on initial behavioral screening, the following seizures were excluded:

i) Seizures in which intact or impaired behavioral responsiveness could not be determined (e.g., no interaction with the patient peri-ictally, or video and audio quality was not adequate).
ii) Electrographic seizures with no clinical manifestations, a.k.a. subclinical electrographic seizures.
iii) Seizures where the only means of determining behavioral responsiveness was by event button pushed without further evaluation.
iv) Cluster seizures during which patients never returned to their baseline level of consciousness between seizures.

This left us with 161 eligible seizures. Subsequently, based on icEEG evaluation, the following seizures were excluded:

i) Seizure onset was not exclusively in the frontal lobe.
ii) Seizures with simultaneous bilateral frontal onset.
iii) Clinical onset preceded the electrical onset (implying true electrographic onset was unknown).

This further left us with 86 eligible seizures.

In our behavioral analysis (see next section), we found that in agreement with previous studies of epilepsy inpatients,^37–39^ testing of responsiveness to questions or commands during seizures was done more reliably than evaluation of patients’ postictal recall of events that happened during seizures. We found that among these eligible seizures that had behavioral testing, 90% were evaluated for ictal responsiveness, whereas only 22% were evaluated postictally for recall of ictal experiences (12% were evaluated for both). Therefore, as has been done for most other studies of impaired consciousness in epilepsy in the inpatient setting ^6,8,11,36,40–47^, we used impaired responsiveness during seizures as the method for evaluating consciousness. This left us with 78 seizures, among which 13 more seizures were excluded due to inconclusive behavioral score, which is defined below in the Behavioral analysis section.

In the end, a total of 65 seizures from 30 patients were used for the final analysis: 18 from Yale, four from Mount Sinai, and eight from NYU. Clinical and demographic information for all included patients can be found in Supplementary Table S1.

### Behavioral analysis

Onset and offset times of all seizures (see Electroencephalography analysis) were provided to a reviewer (ES) who reviewed video of seizures blinded to the EEG recordings. Behavioral analysis proceeded in two stages. In the initial stage of behavioral analysis, the reviewer screened seizures for exclusion based on criteria described in the preceding section. These criteria included lack of behavioral assessment during seizures, subclinical electrographic seizures, and seizure clusters with no behavioral return to baseline between events. In addition, focal to bilateral tonic-clonic seizures were identified based on typical behavioral features, and the time of the onset of generalization was determined behaviorally based on head or eye version, vocalization or asymmetric tonic facial contraction, as in previous studies of FBTC seizures.^48–51^

In the second stage of behavioral analysis, the reviewer performed a detailed assessment of the level of consciousness during all included seizures. This involved the following steps:

1- For each seizure, all stimuli that should have elicited a response in a normal awake individual and their associated responses were documented in detail.
2- Each response was scored as impaired or spared based on the following criteria:
  i) If no response was elicited at all, it was scored as impaired.
  ii) If the only response was that the patient oriented the eyes and/or head toward the stimulus, it was scored as impaired.
  iii) If the patient moved his/her limbs and/or made sounds in response to stimulus in an unintelligible way, the response was scored as impaired. Sounds or movements that were made as a part of typical seizure manifestations were not listed as responses.
  iv) If the patient responded to the stimulus meaningfully and in an appropriate way, the response was scored as spared.
3- Subsequently, the overall behavior during the seizure was determined as follows:
  i) When all responses during the seizure were spared, the seizure was scored as spared.
  ii) When all responses during the seizure were impaired, the seizure was scored as impaired.
  iii) When some of the responses during the seizure were spared but others were impaired, the seizure was scored as inconclusive.
4- Lastly, all seizures were categorized in three groups: i) FA = focal seizures with spared behavioral responses by criteria above; ii) FIA = focal seizures with impaired behavioral responses by criteria above; iii) FBTC = focal to bilateral tonic-clonic seizures, as defined above. Note that all FBTC seizures had impaired behavioral responses.

Seizures with inconclusive behavioral scores were excluded from the analysis as mentioned in the previous section. In addition, if the designated reviewer could not make a clear judgment about a particular stimulus and response, two other reviewers also reviewed the videos (AV and HB). The seizure was included in the final analysis only if the other reviewers could come to a clear decision and the majority of reviewers had the same opinion.

### Anatomic localization of electrode positions

Surgical implants included either a combination of subdural grid, strip, and depth electrodes (24 of 30 patients), or depth electrodes only (six of 30 patients), based on pre-implantation clinical hypothesis. Electrode locations were thus determined clinically and were not standardized. Seventeen out of 30 patients had bilateral implants, five had electrodes only in the left hemisphere and eight had electrodes only in the right hemisphere.

High-resolution MRI scans were performed on all patients from all three centers and later used for electrode localization in each participant:

i) At Yale, post-op CT scans were used to identify electrodes. Then post-op CT scan was co-registered to post-op MRI for each patient, followed by co-registering the post-op MRI to the pre-op MRI, as described previously.^52,53^
ii) At Mount Sinai, the post-implantation CT was co-registered to the pre-operative MRI.
iii) At NYU, electrodes were localized via post-op MRI. Subsequently, the post-implant MRI was co-registered with the pre-implant MR image, as described previously.^53,54^

Electrode contacts for each patient were assigned using regions defined previously,^12^ with the Rolandic region split along the central sulcus into frontal and parietal regions, to obtain the following locations:

i) Ipsilateral frontal – medial, lateral and orbital frontal electrodes in the same hemisphere as seizure onset.
ii) Ipsilateral extra-frontal – occipital, parietal and temporal electrodes in the same hemisphere as seizure onset.
iii) Contralateral frontal – medial, lateral and orbital frontal electrodes in the hemisphere contralateral to seizure onset.
iv) Contralateral extra-frontal – occipital, parietal and temporal electrodes in the hemisphere contralateral to seizure onset.

### Intracranial electroencephalography and video recordings

At Yale, icEEG signals were recorded using either Bio-Logic Systems clinical EEG and video monitoring equipment (Bio-Logic Systems Corp., Mundelein, IL, USA) [prior to 2011] or NATUS/Neuroworks software [after 2011] with sampling rates of 256 and 1024 Hz respectively. At NYU, icEEG signals were acquired using either the BMSI 5000/6000 EEG system (Nicolet Biomedical, Inc., Madison, WI, USA) or NATUS/Neuroworks software with sampling rate of 512 Hz. At Mount Sinai Center, icEEG signals were acquired through NATUS/Neuroworks software with sampling rates of 500, 512, and 1000 Hz.

Signals were recorded relative to a reference electrode chosen by the clinical team to minimize visible noise artifact on the EEG in all three centers. Reference electrodes could include skull pegs affixed to the bone, white matter contacts, mastoid process scalp contacts, or inverted intracranial electrode strips with the insulated side facing the brain surface and the conducting side facing the skull. The NATUS recording system also incorporates a built-in noise cancellation circuit which sampled the reference electrode and injected current of opposite signal back through another nearby electrode.

### Electroencephalography analysis

During icEEG analysis, time of electrographic seizure onset and termination were first identified by expert EEG readers (AV, ES) by visual inspection of the recording. Artifacts were also delineated in the record for each channel by visual inspection and were removed from the analysis. EEG signals were then processed by fast Fourier transform analysis in MATLAB R2018b (Mathworks, Natick, MA, USA) in 1 second (s) non-overlapping data segments for each electrode, and average signal power was calculated within the following frequency bands: delta (0.5 to <4 Hz), theta (4 to <8 Hz), alpha (8 to <13 Hz), beta (13 to <25 Hz) and gamma (25 to ≤50 Hz). We normalized power within each frequency band by calculating percent change in power for each 1 second time segment relative to 30 seconds of baseline power just prior to electrographic seizure onset. Thus, percent change in power was expressed as 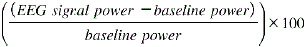 for each frequency band.

As noted above, for calculations of percent change in power, the baseline for all three seizure types was defined as 30 seconds prior to electrographic seizure onset. However, for group analyses across seizures, percent change time course data were aligned to onset (*t* = 0) defined as electrographic seizure onset for FA and FIA seizures, and behavioral onset of generalization for FBTC seizures. We used onset of generalization as *t* = 0 for group data alignment of FBTC seizures rather than electrographic seizure onset because the time between onset and generalization for different FBTC seizures was variable (mean 108.21 s, range 0 to 737 s). Therefore, aligning FBTC seizures to onset of generalization provided more consistent time courses for the FBTC power changes. Percent change in power was then pooled by averaging across electrodes within each of the four analysis regions (Ipsilateral frontal, Ipsilateral extra-frontal, Contralateral frontal, and Contralateral extra-frontal) for each seizure. Finally, mean values and statistics were calculated across seizures for each group of seizures for analysis: FIA, FA, and FBTC.

Group data were plotted as mean (± standard error of mean (SEM)) percent change in power over time in non-overlapping 10 second bins (see Fig. 3C, D; Fig. 5B). For calculation of overall percent power change during seizures versus baseline (see Fig. 3A, B; Fig. 5A; Tables 1, 2) we used the mean for each seizure from onset (defined as *t* = 0 above) to electrographic offset, except for seizures lasting longer than 180 seconds, for which only the first 180 seconds of seizure data were analyzed. For calculating average power changes across all frequency bands (see Fig. 3E, F; Fig. 5C; bottom rows of Tables 1, 2), we first normalized to baseline by calculating percent change from baseline within each frequency band, and then averaged percent change values across frequency bands for each seizure. All above analysis was done using in-house scripts in MATLAB. The result of this analysis was saved in a Microsoft Excel spreadsheet (version 2013) which was further imported to GraphPad Prism 9 (GraphPad Software, San Diego, CA, USA) to generate histogram and time course figures.

To visualize overall power changes anatomically, we generated 3D representations of average percent change in EEG power across all five frequencies for one example of an FA, FIA and FBTC seizure (see Fig. 3E, F; Fig. 5C). This was done using methods described previously.^8,55–57^ Briefly, each electrode contact was localized with the Bioimage Suite software package (www.bioimagesuite.org)^53^ and electrode coordinates were imported into MATLAB.

Reconstruction of the pial surface was based on the patient’s pre-operative T1-weighted high-resolution MRI using the Bioimage Suite software package, and the resulting triangular mesh surface model was also imported into MATLAB. Percent change in power during the seizure from baseline was averaged across the five analyzed frequency bands for each electrode contact, using the methods already described. To display changes on the brain surface, we colored the faces of the triangular mesh based on the power magnitude of the closest electrode contact, applying a linear fade to zero over the radius from 1 to 15 millimeters (mm) surrounding each electrode.

### Statistical Analysis

Statistical tests were performed using SPSS 28 (IBM Corp, Armonk, NY). Percent change in power was analyzed in the ipsilateral frontal regions, and in all other cortical regions combined (see Tables 1,2). We compared percent change in power during FA versus FIA seizures and in FIA versus FBTC seizures using independent two-sample t-tests followed by Holm–Bonferroni correction, with significance assessed at two-tailed *P <* 0.05. All values are presented as mean ± SEM.

## Data availability

All codes generated for data analysis in this study have been deposited in Github (https://github.com) and are available at <<URL will be provided at time of publication>>. All icEEG data for this study are available at DataDryad.org <<URL will be provided at time of publication>>.

## Results

We studied continuous video-intracranial EEG recordings of 65 seizures (24 FA, 27 FIA, and 14 FBTC) in 30 patients. Thirteen patients were female and seventeen were male with a mean age of 27.7 years (range 9-51 years). Twenty-one patients were right-handed, eight were left-handed, and one was ambidextrous. Demographic and clinical information for all included patients can be found in Supplementary Table S1. Seizure type was determined by a blinded video observer who rated patients’ behavior during the seizures (see Methods). Mean duration of FA seizures was 43.2 ± 5.5 s, FIA seizures 56.0 ± 7.6 s, and FBTC seizures 228.6 ± 77.4 s.

### FIA have greater power than FA seizures in wide cortical regions and across frequency bands

We found that FA seizures showed increased icEEG power confined mainly to the frontal lobe of seizure onset, whereas FIA seizures showed increased icEEG power across frequency bands and involving widespread cortical areas both ipsilateral and contralateral to the side of onset. Examples of typical changes are illustrated in representative icEEG recordings for FA (Fig. 1) and FIA seizures (Fig. 2). The example of a FA seizure shows relatively localized ictal findings, with onset as rhythmic spikes within the ipsilateral frontal lobe (Fig. 1A, B) and later spread to part of the ipsilateral parietal lobe (Fig. 1C, D). This contrasts with the FIA seizure example (Fig. 2, same scale as in Fig. 1) which shows early spread beyond the ipsilateral frontal lobe and continued seizure evolution in other bilateral brain regions across all frequency bands. The FIA seizure example begins with onset as low voltage fast in the ipsilateral frontal lobe (Fig. 2A) and then seizure evolution as irregular poly-spikes in ipsilateral frontal lobe as well as propagation with mixed frequencies in widespread brain regions (Fig. 2B-D).

**Figure 1.**
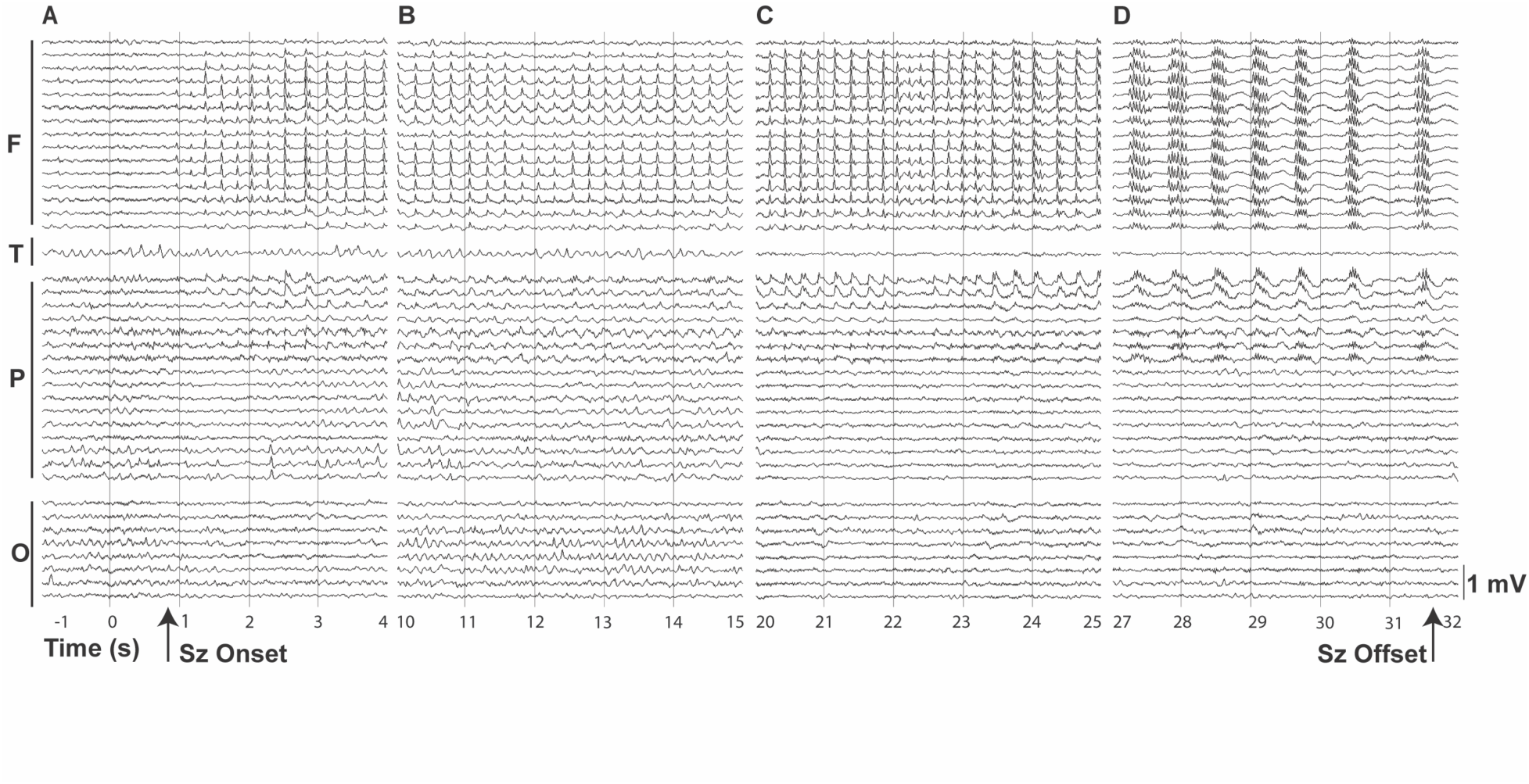
Example of focal aware frontal lobe seizure. Intracranial EEG changes are mainly localized to frontal lobe of onset. **(A)** Seizure (Sz) onset shown by the arrow, with periodic spikes in the frontal electrodes. **(B)** Continuation of seizure as periodic spikes in frontal electrodes and spread of rhythmic activity to some parietal electrodes. **(C)** Evolution in morphology to poly-spike and wave discharges and spread within the frontal and parietal electrodes. **(D)** Seizure offset shown by the arrow in 32nd second as 1 Hz poly-spikes in the frontal and parietal electrodes. Montage is referential to electrode in bone. Calibration bar on the right is 1 millivolt (mV). Horizontal axis shows time in seconds relative to the time interval of seizure onset (0 – 1 s). Illustrative 5 second time epochs are shown from seizure onset to offset. This is a unilateral (left-sided) implantation with a combination of grid, strip, and depth electrodes. Bars on the left show representative electrodes from each lobe. F = frontal; O = occipital; P = parietal; T = temporal.

**Figure 2.**
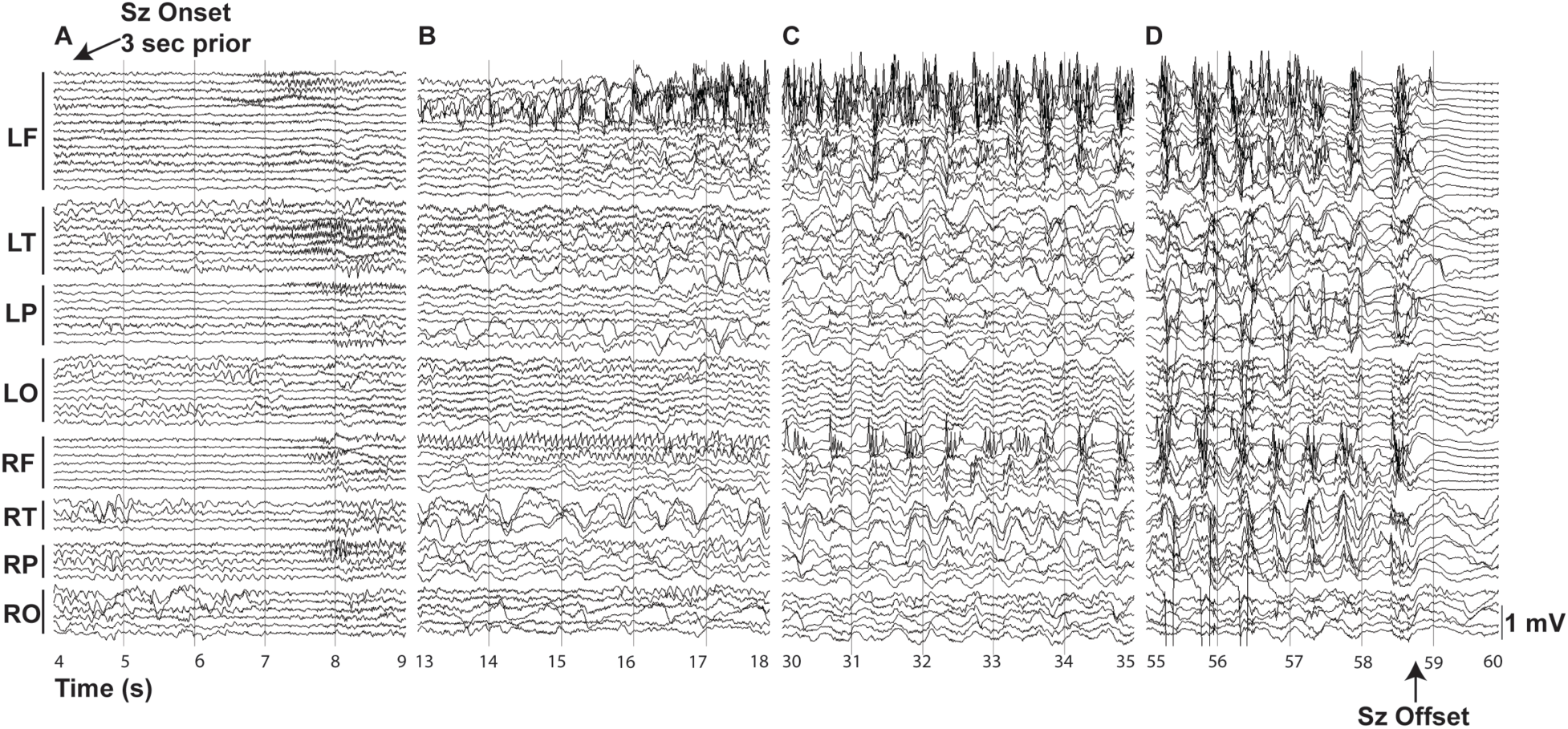
Example of focal impaired awareness frontal lobe seizure. Intracranial EEG changes extend widely beyond frontal lobe of onset. **(A)** Seizure (Sz) onset indicated by arrow with low-voltage fast activity beginning 3 s prior to this time epoch (not visible due to very focal poorly discernible pattern at these settings), evolving as low-voltage fast activity (beta, gamma) in the left frontal contacts with spread to the left temporal, left parietal and right hemispheric contacts. **(B)** Further seizure evolution as irregular poly-spikes with embedded faster frequencies in the ipsilateral (left) frontal electrodes, sharply contoured rhythmic alpha in the contralateral (right) frontal electrodes, and other widespread changes. **(C)** Seizure continues as irregular poly-spikes in bifrontal regions with additional changes in extra-frontal regions bilaterally. **(D)** Widespread 1-2 Hz poly-spike and wave discharges in bilateral hemispheres terminates with seizure offset shown by the arrow in 59th second. Montage is referential to electrode in bone. Calibration bar on the right is 1 millivolt (mV) (same scale as Fig. 1). Horizontal axis shows time in seconds relative to seizure onset. Illustrative 5 second time epochs are shown. This is a bilateral implantation with a combination of grid, strip, and depth electrodes. Bars on the left show representative electrodes from each lobe. LF = left frontal; LO = left occipital; LP = left parietal; LT = left temporal; RF = right frontal; RO = right occipital; RP = right parietal; RT = right temporal.

We next used group data to track electrographic signal changes during both FA and FIA seizures, looking at the common EEG frequency bands (delta, theta, alpha, beta, and gamma) in various brain regions (ipsilateral frontal, ipsilateral extra-frontal, contralateral frontal, and contralateral extra-frontal) (Fig. 3). Summary histograms as well as power time course plots show that FA seizures demonstrated icEEG power increases confined mainly to the frontal lobe of seizure onset, with some spread to the ipsilateral extra-frontal regions (Fig. 3 A, C). In contrast, FIA seizures demonstrated increases in power in widespread regions outside the frontal lobe of seizure onset (Fig. 3 B, D; Ipsilateral Extra-Frontal, Contralateral Frontal, Contralateral Extra-Frontal). These power changes outside the frontal lobe of seizure onset for FIA seizures were seen across multiple frequency bands (Fig. 3 B, D).

Statistical analysis confirmed that FIA seizures had greater icEEG power compared to FA seizures in brain regions outside the frontal lobe of seizure onset (Table 1). We analyzed changes in the frontal lobe ipsilateral to seizure onset, and in all other brain regions combined (Ipsilateral Extra-Frontal, Contralateral Frontal, and Contralateral Extra-Frontal). Cortical power in other brain regions outside the frontal lobe of onset was significantly greater for FIA versus FA seizures (Table 1, right columns). This was true for average power across all frequency bands, as well as for most frequency bands individually. Cortical power outside the frontal lobe of onset across all frequency bands increased by 40.4 ± 7.4% for FIA seizures but only by 10.7 ± 2.6% for FA seizures (*p* < 0.001; Table 1, bottom right entry). In addition, FIA versus FA power was significantly greater for each frequency band individually, with the exception of theta frequency power which did not attain statistical significance (Table 1, right columns). In the frontal lobe ipsilateral to seizure onset, icEEG power was not substantially different for FA versus FIA seizures, except for at higher frequencies where gamma power was significantly greater for FIA than for FA seizures (Table 1, left columns).

To visualize the anatomical distribution of seizure activity more clearly, we created 3D color maps of the average power changes across frequency bands for representative examples of FA and FIA seizures (Fig. 3E, F respectively). These maps show a large increase in the average power across frequencies in all regions that were covered by intracranial electrodes in the FIA seizure (Fig. 3 F). In the example of a FA seizure, the greatest power increases are in the left frontal lobe, which in this case is ipsilateral to seizure onset, with relatively smaller power increases seen in other brain regions (Fig. 3 E).

### FBTC have much greater power than FIA seizures across frequencies in all cortical areas

We have demonstrated that unlike FA seizures, which remain relatively confined to the frontal lobe of onset, FIA seizures exhibit increased icEEG power across frequencies in wide cortical areas, suggesting some degree of seizure propagation throughout the cortex. This raises the question of how frontal lobe FIA seizures differ electrographically from FBTC seizures, which are known to also involve widespread areas of cortex. Therefore, we investigated FBTC seizures that occurred in the same patient group as the FIA seizures, where FBTC seizures were distinguished by typical clinical manifestations including head or eye version, vocalization, tonic facial contraction, and tonic evolving to bilateral clonic limb movements, as in previous studies.^48,49^

We found that FBTC seizures showed much greater icEEG power throughout the cortex and across frequency bands compared to FIA seizures. An example of typical icEEG changes for a frontal lobe onset FBTC is shown in Fig. 4. This seizure shows an initial burst of high amplitude poly-spikes in the ipsilateral frontal lobe (Fig. 4A), followed by rapid bilateral, high amplitude extra-frontal spread (Fig. 4B), evolving to diffuse high amplitude irregular poly-spikes (Fig. 4C) and finally rhythmic poly-spike wave discharges before seizure termination (Fig. 4D). Note that the calibration bar for the FBTC (Fig. 4, lower right) differs from the FA and FIA seizure examples (Figs. 1, 2) illustrating the striking difference in amplitude of the raw EEG waveforms.

**Figure 3.**
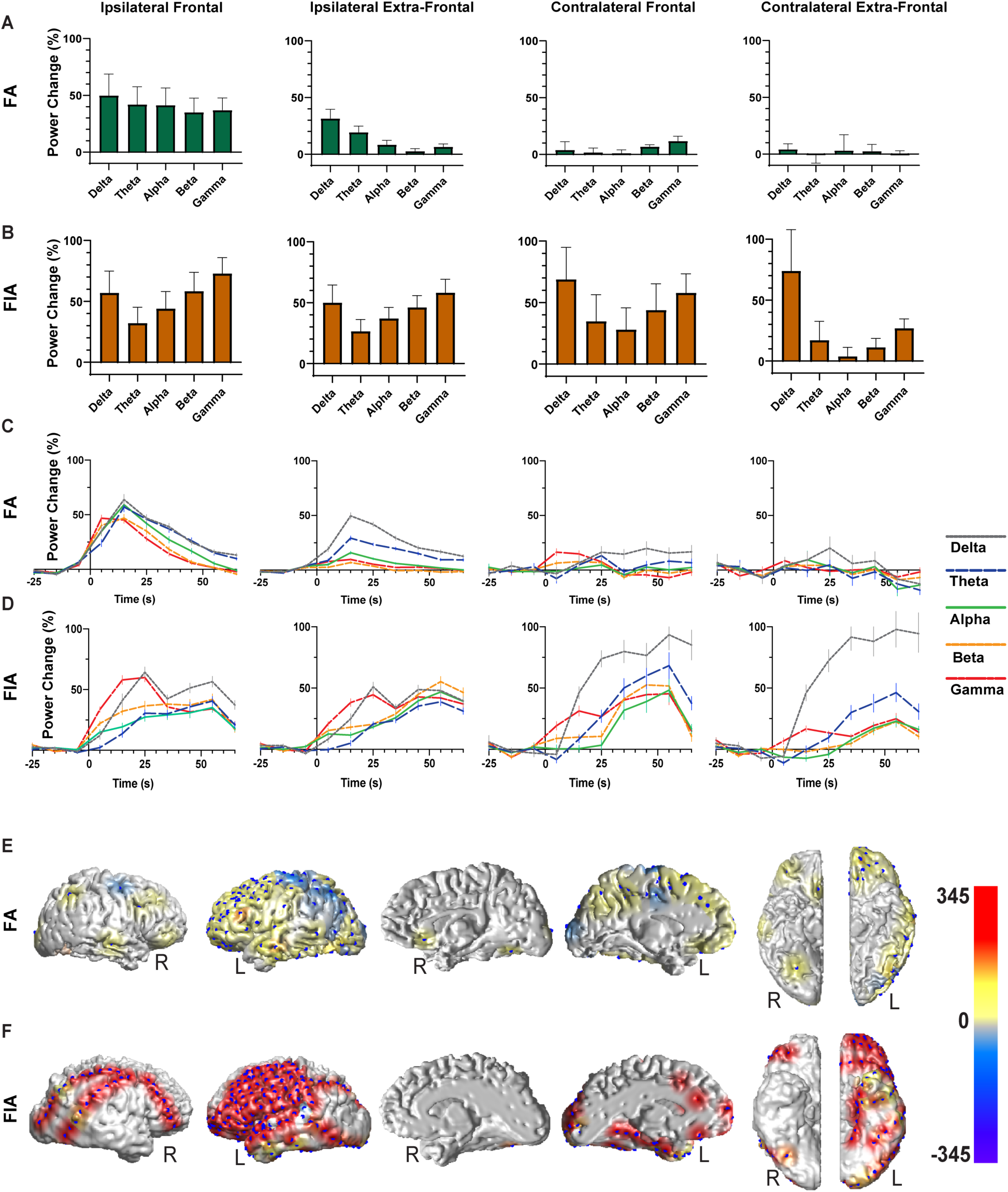
Group analysis and surface maps of icEEG changes during focal aware and focal impaired awareness frontal lobe seizures. **(A, B)** Mean percent change in power in **(A)** focal aware seizures (FA) and **(B)** focal impaired awareness seizures (FIA) compared with 30 seconds pre-seizure baseline. **(C, D)** Time course plots of icEEG percent changes in power during FA seizures **(C)** and FIA seizures **(D)** compared to 30 seconds pre-seizure baseline binned every 10 seconds. Time = 0 is electrographic seizure onset. **(A-D)** Data are from 24 FA seizures in 11 patients and 27 FIA seizures in 16 patients. Error bars are SEM. Delta = 0.5 to <4 Hz; Theta = 4 to <8 Hz; Alpha = 8 to <13 Hz; Beta = 13 to <25 Hz; and Gamma = 25 to ≤50 Hz. **(E, F)** Three-dimensional color maps showing the mean percent changes in power across all five frequency bands versus 30 seconds of pre-seizure baseline in examples of a FA seizure with left frontal onset **(E)** and a FIA seizure with left frontal onset **(F).**

**Figure 4.**
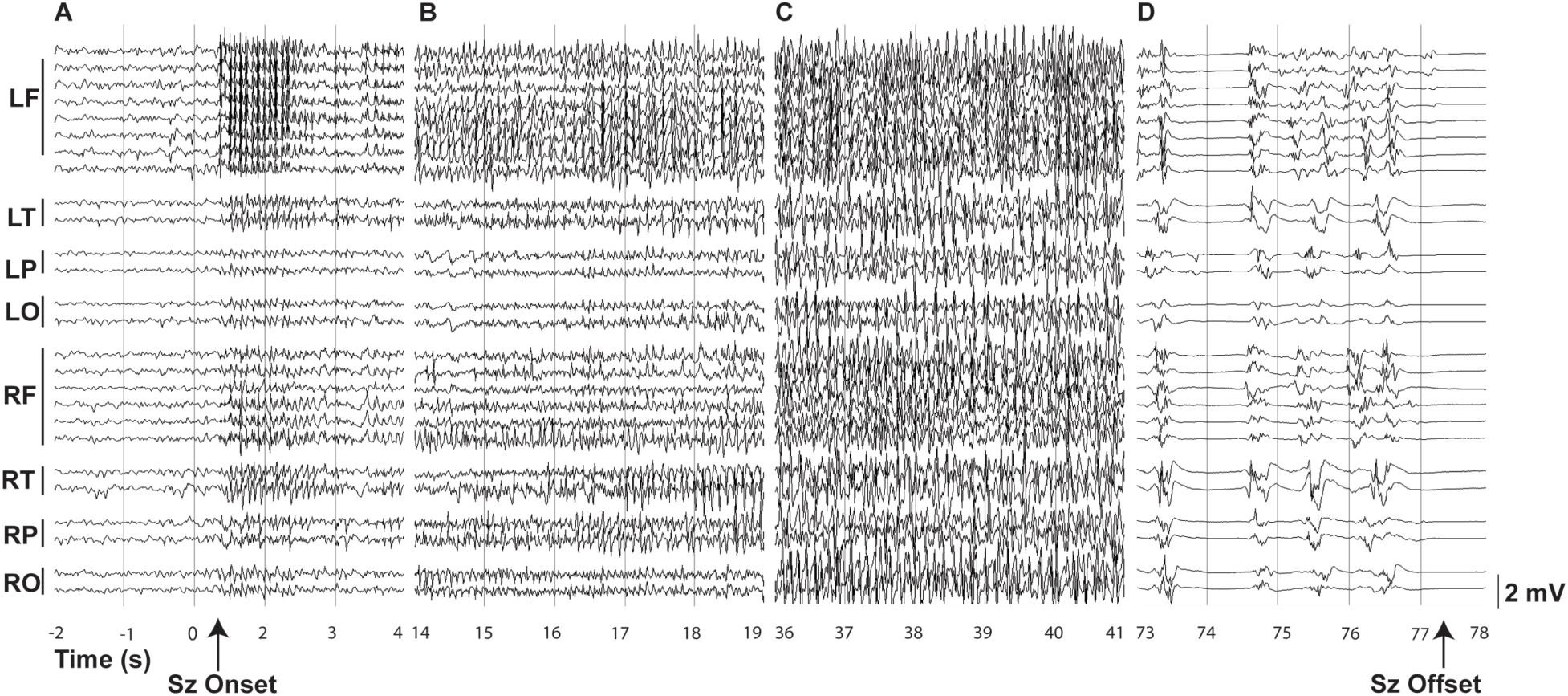
Example of focal to bilateral tonic-clonic seizure of frontal onset. Intracranial EEG changes are widespread and large amplitude. **(A)** Seizure (Sz) onset indicated by arrow, with burst of high-amplitude poly-spikes followed by rapid bilateral and extra-frontal spread. **(B)** Seizure evolution as irregular poly-spikes with embedded faster frequencies in ipsilateral (left) frontal and temporal electrodes with spread to the extra-frontal ipsi and contralateral regions by the end of the panel. **(C)** Tonic phase of the seizure seen on EEG as high-amplitude poly-spikes with admixed mixed frequencies showing widespread bilateral cortical involvement. **(D)** Clonic phase of the seizure shown as irregular ∼1 Hz poly-spike wave discharges with seizure offset shown by the arrow in 78^th^ second. Montage is referential to electrode in bone. Calibration bar on the right is 2 millivolt (mV) (note scale difference from Figs. 1 and 2). Horizontal axis shows time in seconds relative to the time interval of seizure onset (0 – 1 s). Illustrative 5 second time epochs are shown. This is a bilateral implantation with a combination of grid, strips, and depth electrodes. Bars on the left show representative electrodes from each lobe. LF = left frontal; LO = left occipital; LP = left parietal; LT = left temporal; RF = right frontal; RP = right parietal; RO = right occipital; RT = right temporal.

We next used group data to investigate electrographic signal changes during FBTC seizures (Fig. 5). Summary histograms as well as power time course plots show that FBTC seizures demonstrated large amplitude icEEG power increases across cortical regions and frequency bands (Fig. 5 A, B). Statistical analysis confirmed that FBTC seizures had much greater icEEG power compared to FIA seizures in all brain regions and frequency bands (Table 2). For example, icEEG power increase across all frequency bands in the frontal lobe ipsilateral to onset was 52.9 ± 13.4% for FIA and 608.9 ± 104% for FBTC seizures (*p* < 0.001); power across all frequencies in other cortical regions was 40.4 ± 7.4% for FIA and 637.2 ± 107.1% for FBTC seizures (*p* < 0.001; Table 2, bottom row). Similarly, power was also significantly greater for FBTC versus FIA seizures in each individual frequency band, both in the frontal lobe ipsilateral to onset, and in other cortical regions (Table 2).

To illustrate these large magnitude and widespread changes for FBTC seizures, we again created 3D color maps showing the average power changes across frequency bands for an example of a FBTC seizure (Fig. 5C). In this FBTC seizure, there was a large increase in the average power across all regions that were covered by intracranial electrodes, which was considerably higher in scale in comparison to typical FA or FIA seizures (Figs. 3E, F).

**Figure 5.**
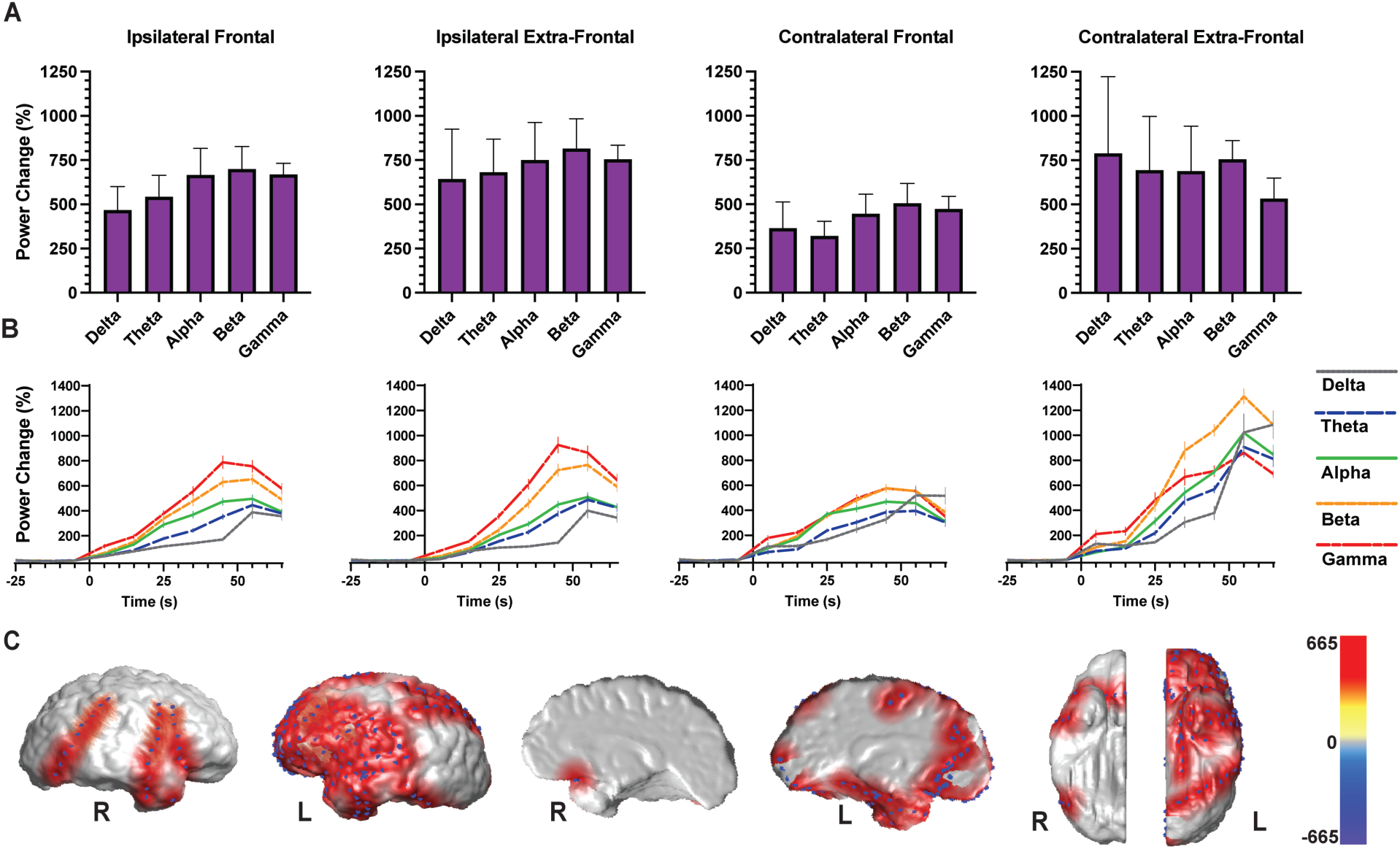
Group analysis and surface maps of icEEG changes during focal to bilateral tonic-clonic seizures of frontal onset. **(A)** Mean percent change in power in focal to bilateral tonic-clonic (FBTC) seizures compared with 30 seconds pre-seizure baseline. Ictal data begin with onset of generalization, however very similar results were obtained if instead ictal data began with electrographic seizure onset (data not shown). **(B)** Time course plots of icEEG percent changes in power during FBTC seizures compared to 30 seconds pre-seizure baseline binned every 10 seconds. Data before Time = 0 are before electrographic seizure onset, data after Time = 0 are after onset of generalization. **(A, B)** Data are from 14 FBTC seizures in 7 patients. Error bars are SEM. Delta = 0.5 to <4 Hz; Theta = 4 to <8 Hz; Alpha = 8 to <13 Hz; Beta = 13 to <25 Hz; and Gamma = 25 to ≤50 Hz. **(C)** Three-dimensional color maps showing the mean percent changes in power across all five frequency bands versus 30 seconds pre-seizure baseline in an example of a FBTC seizure with left frontal onset.

## Discussion

Although there are several studies about physiological distinctions between temporal lobe FIA and FA seizures,^6,8,11,12,58^ there are limited studies focusing on mechanisms of impaired versus spared consciousness in focal FLE, the second most common form of epilepsy. Accordingly, and to elaborate on previous work in understanding the mechanism of impaired consciousness in focal seizures, we have directed our attention to focal FLE. In the present study, we investigated icEEG signals during frontal onset FIA and FA seizures. We found that FIA seizures had greater broadband power increases across frequencies in widespread areas of the cortex whereas FA seizures had relatively localized increases in the frontal lobe of onset. In addition, we found that frontal lobe FIA and FBTC seizures both have widespread icEEG activity increases across frequencies and cortical regions, but the magnitude of the increases is much larger in FBTC seizures.

Consciousness is proposed to depend on normal activity in widespread bilateral cortical networks modulated by subcortical arousal systems. One theory of impaired consciousness in focal epilepsy, the global workspace theory, proposes that widespread abnormal synchrony in cortical and subcortical networks disrupts consciousness during seizures.^7^ Based on this theory, abnormal synchrony has been related to impaired consciousness in TLE,^6,11^ parietal lobe seizures,^40^ as well as in FLE.^36^ A second theory, the network inhibition hypothesis, proposes that inhibited subcortical arousal causes encephalopathy-like cortical delta frequency slow waves and impaired consciousness specifically in TLE.^2,13^ Previously, icEEG work in TLE has shown that slow delta frequency activity increases are significantly larger in bilateral frontoparietal association cortices during temporal lobe seizures with impaired versus spared consciousness.^8,12^ Large amplitude 1– 2 Hz neocortical slow waves observed during temporal lobe FIA seizures closely resemble cortical slowing during other states of unconsciousness such as sleep, coma or encephalopathy.^59,60^ Cortical slow waves during FIA in TLE are associated with widespread frontoparietal decreases in cerebral blood flow correlated with abnormal subcortical activity in patients as well as in animal models.^21,58^ We therefore proposed that in temporal lobe FIA seizures, fast ictal discharges propagate into the contralateral temporal lobe and disrupt the function of midline subcortical arousal systems. This in turn causes the neocortex to enter a depressed state. However, during temporal lobe FA seizures, ictal fast activity remains confined to the seizure initiation region and does not disrupt subcortical arousal systems.^8,13,61^

Interestingly, our present findings show that the mechanisms by which FLE leads to impaired consciousness are different from the above-mentioned mechanisms in TLE. We found that unlike TLE, in frontal lobe FIA seizures, there is an increase in icEEG activity outside the region of seizure onset across a wide range of frequencies, not just in the slow delta frequency range. Based on our findings, instead of indirect effects of seizures on subcortical arousal, frontal lobe seizure activity appears to directly propagate outside the region where it begins focally to involve other widespread neocortices in both hemispheres. The phenomenon of seizure spread throughout the cortex in frontal lobe FIA seizures causes a significantly larger elevation in electrical activity across frequency bands in extra-focal regions during frontal FIA seizures compared to FA seizures. In contrast, frontal lobe FA seizures produce relatively localized increases in icEEG activity across frequency bands confined mainly to the frontal lobe on the unilateral side of seizure onset.

The above observations raised the question whether the widespread icEEG changes in frontal FIA seizures simply can be interpreted as a form of secondary generalization like that seen in FBTC seizures, well known to cause impaired consciousness.^44,48–50^ To address this question, we added a third group categorized as frontal lobe FBTC seizures. The distinction between FIA and FBTC was made by clinical manifestations observed during each seizure. Of note, all seizures in the category of FBTC showed typical impaired consciousness as a part of their behavioral manifestations. Intriguingly, signal properties of these two groups (FIA and FBTC seizures) were substantially different. The elevation in electrical activity across frequency bands was significantly larger by an order of magnitude in both the frontal lobe initiation site and throughout the cortex in FBTC compared to FIA seizures (Figs. 3, 5; Table 2).

These findings suggest a model in which the behavioral severity of seizures, including degree of impaired consciousness, is related to the physiological severity of seizure activity and its impact on widespread regions of cortex. The mechanisms of physiological disruption may vary in different seizure types and can include a combination of factors such as enhanced synchrony per the global neuronal workspace theory, increased cortical slow wave activity per the network inhibition hypothesis in TLE, increased activity across frequencies in FLE, and other activity patterns in different types of seizures. For example, recent work in absence epilepsy has shown that generalized spike wave discharges do not always cause impaired consciousness, but that the degree of behavioral impairment is directly related to the physiological severity of spike-wave EEG amplitude and functional MRI (fMRI) signal changes in widespread corticothalamic networks during seizures.^62–66^ A similar relation between physiological severity and impaired consciousness may also be present in frontal lobe seizures but through different physiological mechanisms. Thus, frontal lobe FA seizures with minimal involvement of widespread cortical areas do not impair consciousness, frontal lobe FIA seizures with widespread cortical involvement across frequencies do impair consciousness, and FTBC seizures with much greater magnitude of ictal activity in widespread cortical areas also cause impaired consciousness. Of note, previous work has shown that the level of impaired behavioral responsiveness is more severe in FBTC seizures than in FIA seizures.^44^

Future work should investigate several open questions not addressed in the present study. For example, despite providing a better understanding of electrophysiological signals associated with decreased consciousness in FLE, our data cannot be used to infer the sequence of events through which abnormal activity spreads throughout the cortex in frontal lobe FIA seizures. There are a few potential circuits which could play role in propagation of seizure activity, among which cortico-cortical and cortico-thalamo-cortical circuits stand out.^67^

Generally, the intralaminar and medial thalamic nuclei form extensive cortico-thalamo-cortical pathways, proposed to modulate synchrony and large-scale integration of information across multiple cortical circuits. Support for this notion include robust connections between intralaminar thalamus and fronto-parietal cortex, playing an important role in arousal regulation,^68^ and diffusion tractography demonstrating extensive interconnections between the thalamic mediodorsal nuclei and frontal cortex.^69,70^ Another thalamic nuclear candidate for cortico-thalamo-cortical propagation is the medial pulvinar given its crucial role for signal coordination in fronto-parietal networks.^71^ Recent studies suggest differential involvement of different thalamic nuclei in focal limbic seizures which may also be relevant for widespread propagation.^19,72^ Additionally, frontal connectivity is certainly compatible with the possibility of seizure spread through cortico-cortical circuits.^73^ In recent decades, corticocortical evoked potentials (CCEPs) have provided extensive evidence that impulses can propagate through direct cortico-cortical white matter fiber pathways.^74–78^ Of course the corpus callosum is also a pathway through which seizures can spread from one hemisphere to the other.^79^ Another important future direction will be more localized investigation of the anatomical regions contributing to impaired consciousness in FLE. Despite investigating a relatively large number of patients and seizures across three centers, and the presence of many cortical electrodes due to the common use of grids and strips rather than stereo-EEG during the time period covered (1997 – 2020), we did not have sufficient electrode coverage across patients to separately study different lobes in each hemisphere. In particular, fewer recordings were available for the hemisphere contralateral to seizure onset especially in extra-frontal regions. Future work with even larger samples across centers should enable investigation of smaller subregions within each hemisphere. Finally, an important question that cannot be addressed with standard clinical icEEG electrodes is how the spread of ictal icEEG activity we observed in FIA seizures across widespread cortical regions and frequencies relates to underlying neuronal firing. For example, it is not known whether the observed widespread cortical icEEG changes are accompanied by changes in neuronal firing, or alternatively local field potential changes related to synaptic inputs without changes in firing. Human unit activity recordings have been used to investigate the distinction between changes in neuronal firing versus local field potential changes ^80^ including studies of ictal unconsciousness,^41^ and this approach could be applied to frontal lobe FIA seizures as well. Finally, just as animal models have been productive for studying underlying neuronal mechanisms of impaired consciousness in absence seizures and TLE,^15,21,65^ an animal model of impaired consciousness in FLE would be very useful for future investigation of fundamental mechanisms.

Impaired consciousness has substantial negative impacts on quality of life of patients with epilepsy.^81,82^ For example, impaired consciousness can lead to motor vehicle accidents, drowning, poor work and school performance, and social stigmatization. Although FLE has not attracted as much attention as TLE due to its less common occurrence, a number of studies of FLE in children have shown that impaired cognitive function and behavior are frequent complications.^83–86^ These comorbidities are correlated with smaller volumes in frontal and extra-frontal regions as well as white matter abnormalities on neuroimaging.^87^ As such, a better appreciation of long-range network effects could help with guiding novel therapeutic strategies to prevent these neocortical consequences and adverse outcomes.

In conclusion, the present study provides quantitative evidence for a broad increase in icEEG activity across frequency bands in widespread regions of cortex associated with impaired consciousness in frontal lobe FIA, whereas FBTC seizures show much larger increases in broadband activity throughout the cortex. These findings contrast with impaired consciousness in focal TLE, where impaired consciousness is associated with widespread cortical slow wave (1-4 Hz) activity. We can speculate that different types of focal seizures produce impaired consciousness by impacting widespread cortical regions but through different mechanisms. Given that cortical dysfunction affects patients’ quality of life, understanding mechanisms underlying this impairment and developing new therapeutic options to prevent it remain important goals for helping people with epilepsy.

## Acknowledgements

We are grateful to the patients who participated in this research.

## Funding

This work was supported by NIH UG3/UH3 NS112826, by the Loughridge Williams Foundation and by the Betsy and Jonathan Blattmachr family.

## Competing interests

The authors report no competing interests.

## Supplementary material

Supplementary material is available at ‘Brain online’.

## Notes

### Competing Interest Statement

The authors have declared no competing interest.

