## Supplementary Table S1 for "Widespread brain activity increases in frontal lobe seizures with impaired consciousness"

*__Supplementary Materials for:__*

Affiliations:
Departments of [1]Neurology, [2]Psychiatry, [3]Neurosurgery and [6]Neuroscience, Yale School of Medicine, New Haven, CT 06520
[4]Department of Neurology, Mount Sinai School of Medicine, New York, NY 10029
[5]Department of Neurology, New York University School of Medicine, New York, NY 10017

**Supplementary Table S1. General characteristics of patient population**

| Pt | Center | Age (yrs) | Sex | H | icEEG Onset | MRI | PET hypometabolism | Ictal SPECT | Surgical intervention | Pathology | Outcome | Follow-up duration |
|---|---|---|---|---|---|---|---|---|---|---|---|---|
| 1 | NYU | 26 | M | R | Left OF | Left sphenoethmoid basal encephalocele | N/A | Left frontal, Left TP | Resection (Left frontal lesionectomy of encephalocele) | Atypical DNET in basal encephalocele; cortical dysplasia + WMH in left frontal basal brain | Engel Class I | 8 years |
| 2 | NYU | 16 | F | R | Right MF, LF | Non-lesional | N/A | N/A | Resection (Right frontal lobectomy) | Cortical dysplasia, white matter neuronal heterotropia | Engel Class I | 6 years |
| 3 | NYU | 25 | F | R | Left MF | Left frontal resection, left genu of CC stroke | N/A | N/A | Staged resection of left mesial superior frontal lobe during icEEG | Prior surgery with cortical dysplasia type I | Engel Class II | 10 years |
| 4 | NYU | 30 | F | R | Left MF | Left anterior cingulate FCD | N/A | N/A | Resection (Left cingulate lesionectomy) | Cortical dysplasia | Engel Class I | 4 years |
| 5 | NYU | 44 | F | L | Right MF, LF | Right mesial frontal, at anterior commisure, FCD | N/A | Left frontal, biparietal, right BG | Resection (Right SMA) + RNS | Cortical dysplasia | Engel Class II | 4 years |
| 6 | NYU | 33 | M | R | Left MF | Prior resection L FP vertex, L posterior fossa arachnoid cyst | N/A | N/A | RNS (Left frontal + Left precentral) | N/A | Engel Class II | 3 years |
| 7 | NYU | 37 | M | L | Left OF, LF | Non-lesional | Bitemporal (L>R) | Normal | Left inferior frontal resection | Cortical dysplasia | Engel Class I | 3 years |
| 8 | NYU | 39 | M | R | Right LF | Bi-occipital (R>L) MCD | R > L posterior | Right temporo-parieto-occipital | Right inferior frontal corticectomy | Reactive gliosis | Engel Class I | 1 year |
| 9 | Yale | 22 | M | AD | Left LF, OF | Left frontal FCD | L inferior frontal | N/A | RNS (Left inferior frontal + anterolateral temporal) | N/A | Engel Class III | 7 years |
| 10 | Yale | 34 | M | R | Left LF | Non-lesional | R temporal | N/A | None (declined) | N/A | N/A | N/A |
| 11 | Yale | 40 | M | R | Left MF | Left inferior temporal possible FCD | Negative | N/A | RNS (Left medial frontal + Left mid frontal) | N/A | Engel Class II | 3 years |
| 12 | Yale | 18 | M | L | Right LF | Right frontal FCD | Right frontal | N/A | Right lateral frontal resection | Cortical dysplasia | Engel Class I | 3 years |
| 13 | Yale | 15 | F | L | Left LF, OF | Left hemisphere cyst, parieto-occipital predominant | L non-localizing | Left frontal | Left anterior cingulate resection + corpus callosotomy | Focal mildly increased neurons in subcortical white matter, subpial and intraparenchymal reactive gliosis | Engel Class I | 3 years |
| 14 | Yale | 34 | M | L | Left LF | Non-lesional | L fronto-parieto-temporal | N/A | Left frontal resection | Normal | Engel Class I | 1 year |

| | | | | | | | | | | | | |
|---|---|---|---|---|---|---|---|---|---|---|---|---|
| 15 | Yale | 23 | M | R | Right OF | Non-lesional (Right parietal DVA) | R fronto-parietal | Right temporal, Left posterior | Right frontal resection | Right polymicrogyria | Engel Class I | 16 years |
| 16 | Yale | 27 | F | L | Left MF | Non-lesional | R Temporal | Left frontal | Left frontal (SMA) resection | Normal | Engel Class I | 5 years |
| 17 | Yale | 27 | M | L | Left OF | Non-lesional | L temporal | Negative | Left frontal pole resection | Chronic gliosis, scattered subcortical neurons | Engel Class IV | 11 years |
| 18 | Yale | 18 | F | R | Left LF | Non-lesional | L fronto-parietal | Left fronto-parietal | RNS (Left frontal +Left parietal) | N/A | Engel Class II | 3 years |
| 19 | Yale | 9 | M | R | Right LF | Right frontal FCD | R parietal | Right parietal | Right frontal resection + MST | Cortical dysplasia | Engel Class III | 11 years |
| 20 | Yale | 51 | F | R | Right LF | Right frontal AVM resection cavity | R temporal | Right temporal | Right frontal resection | Reactive gliosis, chronic ischemia | Engel Class IV | 7 years |
| 21 | Yale | 29 | F | R | Right LF | Right superior frontal gyrus thickening | R frontal | Right temporal, Left posterior | Right frontal resection | Normal | Engel Class I | 11 years |
| 22 | Yale | 41 | F | R | Right LF | Non-lesional | Negative | N/A | Right frontal (Primary motor cortex) MST | N/A | Engel Class III | 11 years |
| 23 | Yale | 15 | M | R | Right OF, MF | Non-lesional | R temporal | Negative | Right frontal resection (OF, MF) | Normal | Engel Class IV | 16 years |
| 24 | Yale | 29 | F | R | Right OF | Non-lesional | R hemisphere | N/A | Right frontal resection + MST | Normal | Engel Class III | 10 years |
| 25 | Yale | 13 | M | R | Right LF | Right SFG thickening | Negative | N/A | Right frontal resection | Normal | Engel Class I | 13 years |
| 26 | Yale | 20 | M | R | Left MF, LF | Non-lesional | Negative | N/A | Left frontal resection | Normal | Engel Class II | 12 years |
| 27 | Sinai | 16 | M | L | Left LF | Non-lesional | N/A | N/A | RNS (Left frontal SMA/premotor) | N/A | Engel Class III | 3 years |
| 28 | Sinai | 27 | M | R | Left LF, MF | Right frontal encephalomalacia | N/A | N/A | Right frontal disconnection | Right frontal gliosis (chronic infarct) | Engel Class I | 2 years |
| 29 | Sinai | 34 | M | R | Right LF | Right frontal non-enhancing tumor | N/A | N/A | Bilateral frontal (FEF) RNS | Right frontal anaplastic astrocytoma (WHO Classs III) | Engel Class III | 2 years |
| 30 | Sinai | 38 | F | R | Right LF | Non-lesional | N/A | N/A | Right frontal resection | Right frontal anaplastic astrocytoma (WHO Classs II) | Engel Class III | 5 years |

AD = Ambidextrous; AVM = Arterio-venous malformation; BG = Basal ganglia; DNET = Dysembryblastic neuro-epithelial tumor; DVA = Dural venous anomaly; F = Female; FCD = Focal cortical dysplasia; FEF = Frontal eye fields; FP = Frontoparietal; H = Handedness; icEEG = Intracranial electroencephalography; L = Left; LF = Lateral frontal; M = Male; MCD = Malformation of cortical development; MF = Medial frontal; MRI = Magnetic resonance imaging; MST = multiple subpial transection; N/A = Not available; NYU = New York University School of Medicine; OF = Orbitofrontal; PET = Positron emission tomography; Pt = Patient number; R = Right; RNS = Responsive neurostimulation; SFG = Superior frontal gyrus; Sinai = Mount Sinai School of Medicine; SMA = Supplementary motor area; SPECT = Single photon emission computed tomography; TP = Temporoparietal; WHO = World health organization; WMH = White matter hyperintensities; Yale = Yale School of Medicine; yrs = years.
